# Greater benefits of assisted gene flow in F2 vs F1 progeny at the cold edge of a species’ range

**DOI:** 10.64898/2026.03.14.711431

**Authors:** Brandon Hendrickson, Megan L. DeMarche, Drew Maraglia, Oscar Gonzalez, Kevin J. Rice, Sharon Y. Strauss, Jason P. Sexton

## Abstract

Gene flow to marginal populations at a species’ range edge can facilitate rapid adaptation by increasing genetic diversity, reducing inbreeding depression, and introducing novel alleles. In highly inbred populations, hybrid vigor is often observed in the first generation (F1), but hybrid breakdown may diminish fitness in subsequent generations. Thus, benefits of gene flow may be overestimated when only F1 performance is assessed. We tested whether gene flow among populations of the annual plant *Erythranthe laciniata* (A. Gray) G.L. Nesom, from similar and contrasting environments, confers persistent fitness advantages across F1 and F2 generations at the high-elevation edge of its range in the California Sierra Nevada. Gene flow was experimentally introduced through pollen transfer between cold-edge populations, between cold edge and central populations, and within local cold edge populations, and compared to self-fertilized offspring, the predominant mating strategy of *E. laciniata*. For F1 progeny, we measured morphological, phenological, and fitness traits in a common garden located near the cold-climate range limit during 2008-2009, a relatively average year, and for F2 progeny in 2009-2010, a relatively wet year. Although F1 crosses showed no initial performance advantage measured in the previous year, F2 progeny from center-to-edge and edge-to-edge crosses significantly outperformed selfed and locally outcrossed lines in fruit mass, total pedicels, biomass, and height. Our findings demonstrate that gene flow can confer long-term fitness benefits, especially among populations adapted to similar selective pressures, and highlight the potential of assisted gene flow to bolster or rescue peripheral populations facing climate change.

**SIGNIFICANCE STATEMENT:** Species living at the edges of their geographic ranges often have small, isolated populations with limited genetic diversity, which can restrict their ability to adapt to environmental change. Gene flow from other populations may increase adaptive potential, but its long-term consequences remain uncertain because most studies evaluate only first-generation hybrids. Using experimental crosses in the mountain wildflower *Erythranthe laciniata*, we show that gene flow can produce stronger fitness benefits in second-generation hybrids than in the first generation at a high-elevation range edge. These results suggest that recombination among populations can generate advantageous genetic combinations that emerge over multiple generations. Our findings highlight the potential for assisted gene flow to enhance adaptation and persistence of range-edge populations under climate change.

## INTRODUCTION

Peripheral populations often occupy marginal environments where population sizes are small and connectivity is limited, increasing the influence of inbreeding, genetic drift, and demographic stochasticity (1–3). These processes reduce standing genetic variation and constrain adaptive responses to environmental change (4–6). Such effects are frequently pronounced at leading (cold or high-elevation) range edges shaped by post-glacial colonization, where serial founder events reduce genetic diversity and elevate mutational load (7–10), potentially limiting further range expansion (11). Although genetic diversity within peripheral populations is often reduced, range-edge populations are not evolutionarily trivial. They frequently exhibit strong differentiation among populations and may harbor unique genetic variants, contributing disproportionately to species-wide genetic diversity (5, 8, 12). Genetic exchange among such populations may therefore play an important role in shaping fitness and evolutionary potential at range margins.

Outcrossing among populations represents a key evolutionary process influencing fitness in small or isolated populations. Even low levels of gene exchange can increase heterozygosity, alleviate inbreeding depression, and restore genetic variation lost to drift (13–17). Range-edge populations often experience similar climatic regimes and selective pressures but differ genetically due to drift, founder effects, and independent accumulation of mutational load. Outcrossing among such climatically matched yet genetically differentiated populations may therefore introduce favorable alleles and genetic variation while avoiding the risks associated with environmentally mismatched gene flow (18–23). A common short-term consequence of outcrossing is heterosis, in which F1 hybrids outperform parental genotypes due to masking of deleterious recessive alleles (16, 24–29). Recombination among divergent populations can also generate novel genetic combinations, including transgressive phenotypes that may enhance performance in stressful or marginal environments (26, 30–32). Nevertheless, hybridization can also impose costs. Genetic divergence among populations can generate epistatic incompatibilities that reduce hybrid performance, particularly in later generations (33), and hybrid fitness often declines with increasing divergence between populations (34, 35). Maladaptive effects have been observed in crosses between habitat types (34, 35) and across gradients (36, 37), though a growing number of studies find gene flow between climatically similar, yet distant peripheral populations can increase fitness (20, 38–40). Furthermore, elevated fitness in the F1 generation does not necessarily predict longer-term outcomes, as heterozygosity can mask deleterious interactions that reemerge in the F2 generation (41, 42). Alternatively, favorable recombinant genotypes may persist or increase across generations. Evaluating the evolutionary consequences of outcrossing therefore requires explicit multigenerational tests.

A previous experiment at the low-elevation (warm) range boundary of *Erythranthe laciniata* tested how outcrossing among populations influenced F1 fitness when offspring were grown at the rear edge of the species’ distribution (20). That study showed that fitness responses depended on the geographic and climatic origin of pollen donors and demonstrated that crosses between range-edge populations could substantially increase F1 fitness when grown at the low-elevation boundary. These results highlighted the potential importance of edge-to-edge outcrossing among climatically similar but genetically differentiated populations. Here, we extend this work by testing whether the fitness effects of outcrossing observed at the warm, low-elevation boundary generalize to the cool, high-elevation leading range edge and persist beyond the F1 generation. Using the same species and experimental framework, we sampled populations along the same two elevational transects, but focusing on high-elevation range limit and central populations. We generated crosses between focal high-elevation populations and multiple source populations, including other high-elevation range-edge populations, intermediate lower-elevation populations, and geographically central populations (Table 1). F1 and F2 offspring, along with selfed controls, were grown in a high-elevation common garden across two climatically contrasting years.

**Table 1.** Population information for sires and dams along two transects. The HE dam population (A transect) was crossed with BC, BP, ME, and ML whereas the ML dam population (B transect) was crossed with TN, SC1, ME, and ML. Mean annual temperature (MAT) and precipitation of the wettest quarter (WQPPTN) bioclim variables were collected at a resolution of 0.5 arc-minutes (∼1km^2^ grid size) using the raster R package. Pairwise genetic distance estimates between populations are graph distances estimated from simple sequence repeat (SSR) marker variation reported Sexton et al. 2016. Elevation of each population was collected at a resolution of 0.5 arc-minutes using the elevatr R package (Hollister et al. 2023). Habitat type categories are based on Storer et al. (2004, pp. 20-22).

| Cross Type | Transect - Dam | Sire Population (Lat., Long.) | Genetic distance | MAT (C°) | WQPPTN (mm) | Elevation | Habitat Type |
| --- | --- | --- | --- | --- | --- | --- | --- |
| Local | 1 - HE | HE (37.3562, -118.8608) | NA | 1.4 | 284 | 3095 | Subalpine |
| Close Center | 1 - HE | BC (37.358, -118.8806) | 36.01 | 1.4 | 284 | 2774 | Subalpine |
| Far Center | 1 - HE | BP (37.2384, -119.2598) | 17.54 | 10.4 | 437 | 2473 | Upper Montaine |
| Close Edge | 1 - HE | ME (37.6966, -119.0919) | 46.25 | 2.1 | 281 | 3049 | Subalpine |
| Far Edge | 1 - HE | ML (37.8402, -119.4921) | 31.72 | 3.3 | 345 | 2774 | Subalpine |
| Local | 2 - ML | ML (37.8402, -119.4921) | NA | 3.3 | 345 | 2774 | Subalpine |
| Close Center | 2 - ML | TN (37.8363, -119.4556) | 33.21 | 3.3 | 345 | 2500 | Subalpine |
| Far Center | 2 - ML | SC1 (37.77955, -119.53398) | 42.71 | 5.1 | 383 | 2165 | Upper Montaine |
| Close Edge | 2 - ML | ME (37.6966, -119.0919) | 16.71 | 2.1 | 281 | 3049 | Subalpine |

## RESULTS

Survival to flowering was high in both generations (F1: 80.7%, F2: 73.0%) and did not differ significantly among cross types, transects, or the interaction (Table 2; Fig. 2). Fruit mass was not significantly influenced by cross type in the F1 generation, as crosses did not exceed selfed controls. In contrast, fruit mass in the F2 generation was significantly influenced by cross type (Table 2), with no effects of transect or interactions detected. In the F2 generation, far edge-to-edge, far center-to-edge, and close edge-to-edge crosses all produced significantly greater fruit mass than selfed progeny (Fig. 2; Suppl. Table 2). Model comparisons of Bayesian hurdle models integrating effects across both survival and fruit mass simultaneously (lifetime fitness) (using WAIC; Suppl. Table 3) indicated that overall treatment effects on combined survival and fruit mass were limited for both F1 and F2 plants (Suppl. Table 3).

**Table 2:**
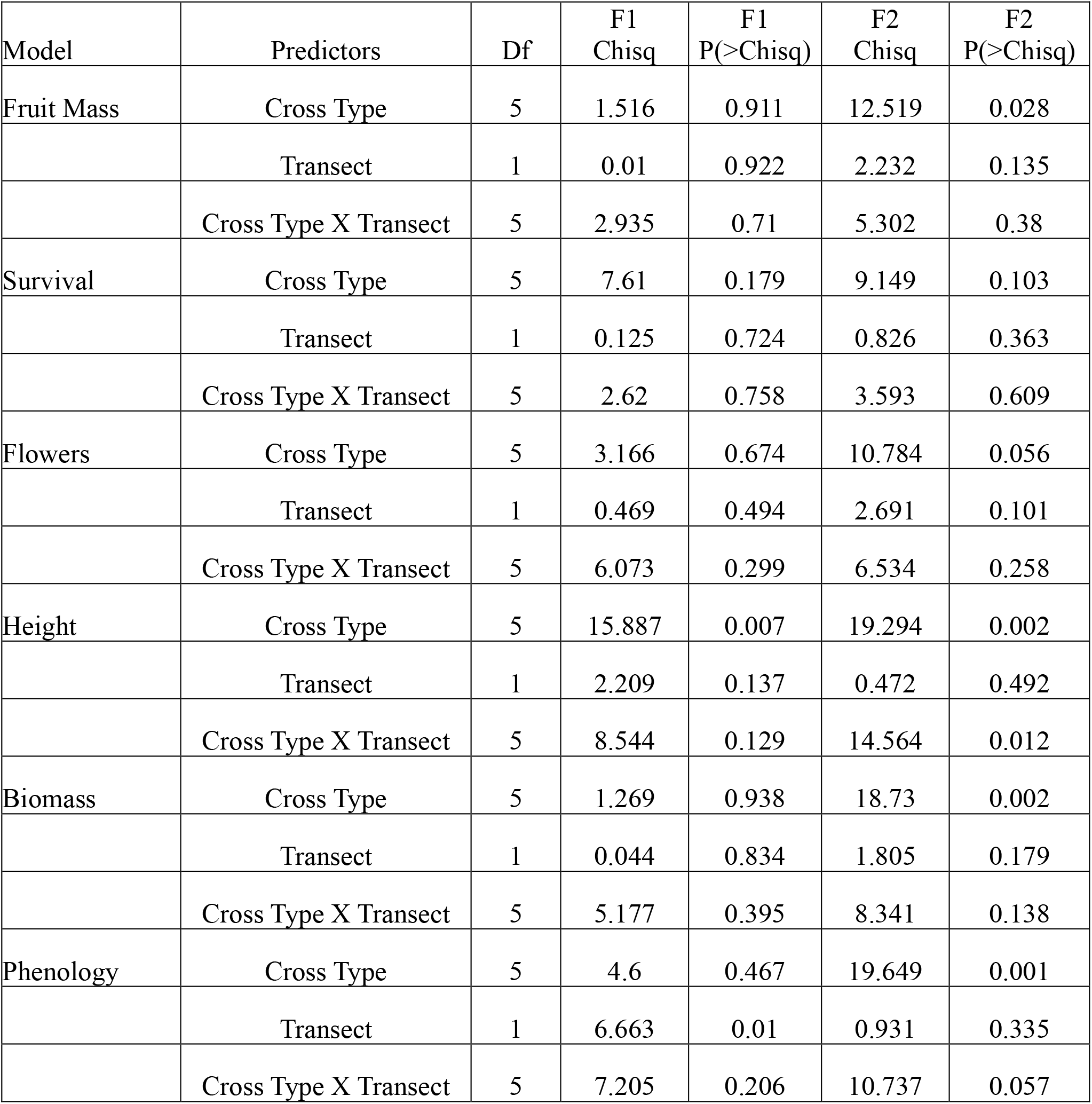
Generalized linear models of F1 and F2 fitness, morphological, and phenology traits at the high-edge range limit of *Erythranthe laciniata*.

Cross type significantly influenced height in both generation years, and total pedicels and biomass in the F2 generation year (Fig. 2; Fig. 3). In the F2 generation year, far edge-to-edge and far center-to-edge crosses were taller and had greater biomass than selfed progeny –translating into significantly greater fruit mass, whereas close edge-to-edge crosses showed significant gains in fruit mass but nonsignificant increases in biomass and height. In the F1 generation year, far edge-to-edge and far center-to-edge progeny were significantly taller than selfed controls but did not produce greater fruit mass (Fig. 2; Suppl. Table 2).

A sub analysis of cross types within transects revealed that the effects of gene flow also depended on transect of origin, a result that has important conservation implications for targeted gene flow efforts. Although the overall interaction between cross type and transect was not statistically significant in the omnibus test, significant effects emerged in planned pairwise contrasts within each transect — a pattern expected when cross type effects are consistent in direction but vary in magnitude across transects, producing localized differences detectable at the transect level but insufficient to drive a significant global interaction term. Far edge-to-edge and close edge-to-edge crosses sired by the away population (i.e., ML) had greater fruit mass, height, and number of total pedicels than selfed progeny from the ML population (Fig. 3; Suppl. Table 5). In addition, home-sired far edge-to-edge progeny showed significant biomass gains and marginal increases in fruit mass (p = 0.061) and height (p = 0.054) compared to selfed controls. Far center-to-edge crosses also differed by transect, with home-sired progeny displaying more advanced phenology than those sired by away populations (Suppl. Table 5).

Phenotypic selection analyses relating fruit production to traits revealed strong directional selection in the F2 generation for greater biomass, taller height, and earlier phenology (Fig. 4). Positive selection on phenology and biomass, but not height, was detected in the F1 generation year and was weaker than in the F2 generation year (Fig. 4). The strength of selection (β) acting on height and biomass was substantially amplified in the F2 generation year relative to the F1 year, with increases of more than ten-fold and nearly three -fold, respectively. Moreover, these traits accounted for far more variation in fruit mass (R^2^) in the F2 than in the F1. Notably, phenology accounted for only a small proportion of fruit mass variation, 1.4% in the F1 year and 1.2% in the F2 year, and the incremental gains in fruit mass associated with shifts in phenology were markedly lower than those attributable to height or biomass. However, positive selection on earlier flowering was observed in both the F1 and F2 generations to an equal degree.

## DISCUSSION

We found beneficial gene flow effects at the high-elevation leading edge between climatically distinct populations (from far away populations occupying similar high-elevation environments and from central areas of the range) that carried through to the F2 generation, suggesting increased adaptive genetic variation through admixture. Gene flow among populations occupying different habitats—such as central versus peripheral or climatically divergent populations—can have heterogeneous effects on hybrid fitness. Classical theory predicts that long-distance or environmentally mismatched gene flow may introduce maladaptive alleles and impede local adaptation (43–46). Empirical support for this view exists: for example, center-to-warm-edge gene flow in *Erythranthe laciniata* resulted in reduced F1 performance (compared to locally outcrossed plants) in warm, dry environments, even while improving emergence relative to selfed local progeny, likely through partial relief of inbreeding depression (20). However, this framework does not fully capture the range of outcomes observed in nature. Gene flow across environmental gradients can also enhance genetic variation, introduce beneficial alleles, and generate hybrid vigor, particularly under environmental change (16, 47). Indeed, highly vigorous F1 hybrids have been reported from crosses between low- and high-elevation populations of *Clarkia xantiana* in the Sierra Nevada (48). Consistent with these broader patterns, we detected no evidence of outbreeding depression. Instead, several crosses between climatically distinct populations exhibited elevated fitness, suggesting that gene flow at the leading edge can be beneficial under certain demographic and environmental contexts. This is another study that rejects the hypothesis that asymmetric gene flow from the center maintains range limits (as in (49)) as well as highlighting the fitness benefits of edge-to-edge gene flow in the context of leading-edge populations.

Hybrid vigor is most commonly expressed in F1 progeny derived from genetically differentiated populations with a history of inbreeding or bottlenecks (20, 50, 51). Inbreeding depression arises through the expression of deleterious recessive alleles or the loss of favorable heterozygotes, and it can reduce both fitness and adaptive potential by depleting additive genetic variance (52–55). Although purging of deleterious alleles is possible in highly selfing populations (56, 57), gene flow remains an important mechanism for masking genetic load and restoring variation (58–60). Surprisingly, we detected neither strong inbreeding depression in high-elevation populations nor pronounced heterosis in F1 hybrids. This pattern is consistent with theoretical expectations for relatively large or demographically stable populations, where mutational load may already be reduced (29, 57, 61). Moreover, these findings support those of Shay et al. 2026, who found that high-elevation populations of this plant species have high expression of adaptive differentiation. Of note, both of the far away gene flow donor populations that produced the greatest fitness effects observed in the F2 generation (ML at the high edge and BP from the range center) are relatively small populations (ML = “C-High”; BP = “A-4” in Table 1 in (62)), and smaller than the home population in this study, HE, demonstrating the potential of small populations to be sources of highly beneficial gene flow.

Several fitness advantages emerged in the F2 generation, particularly for fruit mass, suggesting that recombination rather than simple dominance effects underlies the benefits of outcrossing at the leading edge. Increased F2 fitness may reflect the introgression of alleles better suited to contemporary conditions or the formation of novel, adaptive gene combinations. Importantly, models predicting maladaptation from central-to-edge gene flow typically assume static environments (45, 63, 64). High-elevation *E. laciniata* populations are locally adapted to cold conditions (65), yet under ongoing warming, gene flow from warmer populations may enhance fitness rather than erode it (21). Consistent with this idea, F2 hybrids derived from moderate- and warm-climate populations exhibited increased biomass, height, and fruit mass when grown at high elevation, supporting the potential utility of assisted gene flow under climate change.

Gene flow among climatically similar but genetically differentiated edge populations also emerged as a strong driver of fitness gains. Parallel adaptation to similar environments can result in the fixation of distinct adaptive variants across populations (66–68), which may recombine through hybridization to generate transgressive phenotypes (30). Such effects are expected to be particularly pronounced in highly selfing species like *E. laciniata*, where intraspecific crosses between inbred populations can release hidden variation. Consistent with this expectation, edge-to-edge crosses increased fitness at both low- and high-elevation range margins ((20); this study). Moreover, F2 hybrids from edge-to-edge crosses exhibited significantly greater biomass, height, total pedicels, and fruit mass than selfed controls. These results support the hypothesis that gene flow between similarly adapted populations can facilitate movement across rugged fitness landscapes by enabling access to alternative adaptive peaks (69–71) while avoiding the maladaptive effects often associated with environmentally mismatched gene flow. Our results underscore the need for nuanced, population-informed strategies in conservation and assisted gene flow and also the reality that beneficial gene flow can come from many sources (72, 73).

The delayed expression of fitness benefits until the F2 generation further suggests that recombination, purging, or both play key roles. Hybridization may expose deleterious alleles in additive traits, allowing them to be rapidly eliminated in subsequent generations (74–76). Trait-specific responses also indicate that genetic architecture matters: although fruit mass and reproductive traits improved only in the F2 generation year, height increased in both the F1 generation and F2 generation for certain cross types, implying different sensitivities to dominance, epistasis, additive genetic variance, or the environment. Environmental context likely modulated these outcomes. The F2 generation was grown in a cooler, wetter year that may better approximate high-elevation conditions with greater snowpack. This may partially explain why some cold edge-to-edge crosses performed better in the F2 than in the F1.

Overall, our findings reinforce the idea that gene flow can enhance fitness at range edges (40), but that its benefits depend critically on climatic similarity, population history, and generation. At the leading, high-elevation edge of *E. laciniata*, beneficial gene flow may originate from both climatically similar edge populations and from warmer or wetter populations likely to resemble contemporary and future conditions. Although we could not explicitly test the relationship between genetic distance and fitness, we observed benefits across both relatively low and high differentiation levels (Table 1). Prior work suggests that genetic isolation by environment (climate), rather than geographic distance, structures genetic variation in this species (62), and that climatically matched but genetically distinct populations often perform best near range limits (65).

Finally, our results highlight the benefits of extending beyond the F1 generation when evaluating the evolutionary consequences of gene flow, particularly in highly selfing species. F1 heterosis may overestimate short-term benefits while obscuring longer-term dynamics driven by recombination and selection. To our knowledge, this study provides the first multigenerational experimental test of fitness, morphology, and phenology at a leading range edge under field conditions. Together, our findings emphasize the adaptive potential of both edge-to-edge and center-to-edge gene flow at high elevations and underscore the need for population-informed approaches to conservation and assisted gene flow in a changing climate.

## METHODS & MATERIALS

### Sample Populations

*Erythranthe laciniata* seeds were sampled along two widely spaced geographic transects following the Sierran elevation gradient (Figure 1; Table S1). Additionally, a high-edge population (ME) between transects was sampled to serve as a pollen donor of intermediate distance to two other edge populations. These transects were chosen to capture replicate gradients and were part of different watershed areas (Merced River and North Fork of the San Joaquin River, versus South Fork of the San Joaquin River) occupied by this creek- or seep-dependent species. Range limits were defined as areas supporting the highest elevation populations along the elevation gradient. The range limit for each transect was verified by an exhaustive search within potentially suitable habitats at higher elevations. Monthly temperature and precipitation from 1981 to 2010 were collected using the prism R package (77) and the thirty-year climate average was calculated (Fig. 1). Annual averages of mean, maximum, and minimum temperature, as well as accumulated precipitation, were calculated for the two experimental years, 2008-2009 and 2009-2010 starting from October 1st to September 31st.

**Figure 1.**
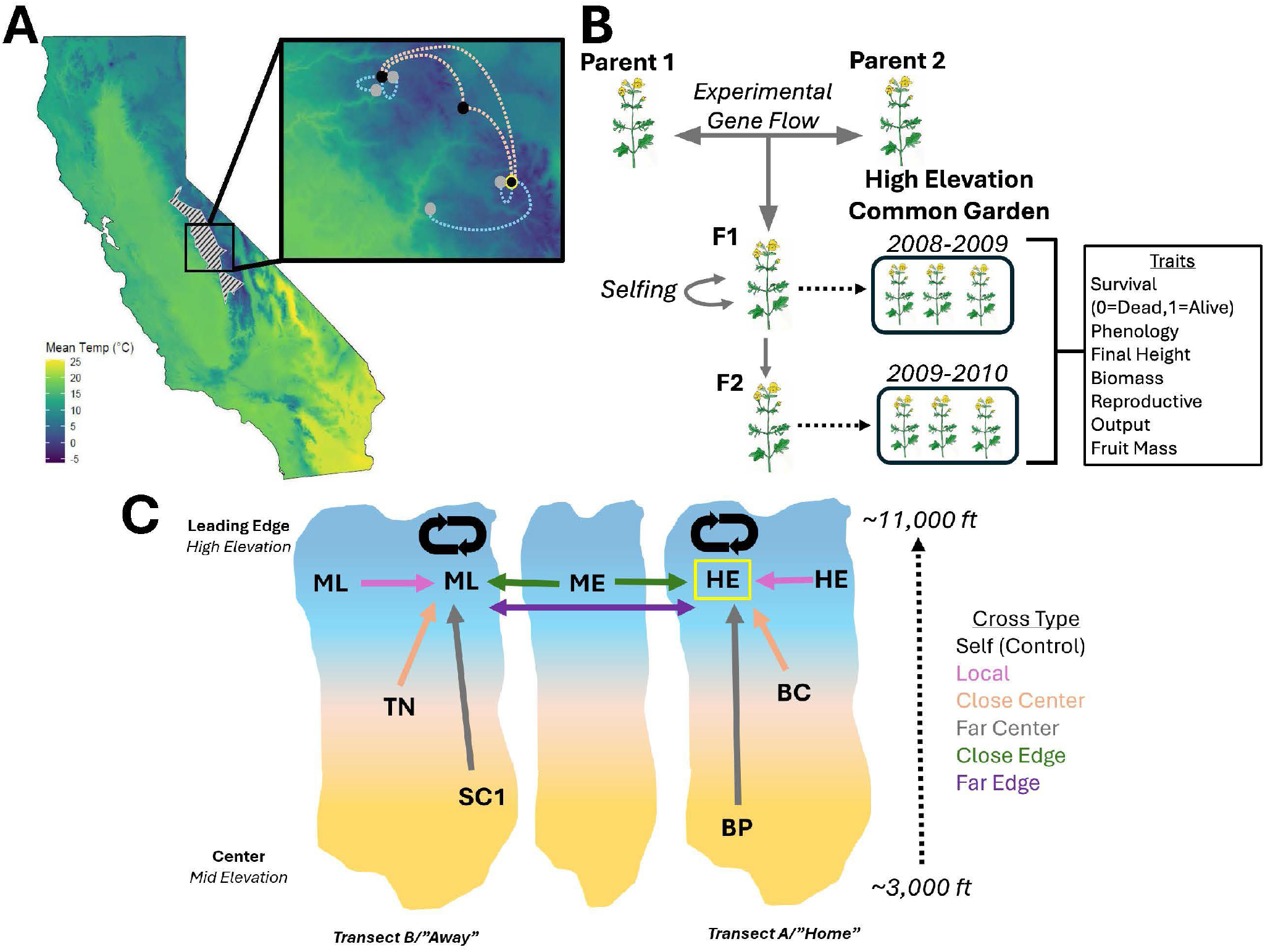
(A) Climate map of California showing the convex hull of the *Erythranthe laciniata* range. The inset (upper right) zooms in on the study region and the seven experimental populations. Populations from the home transect (HE, BC, BP) and the away transect (ML, TN, SC1) are distinguished by color, with peripheral populations in black and central populations in grey for each transect. Dotted lines connecting populations represent experimental gene flow, with light blue being within transect crosses and orange lines representing edge to edge gene flow. The location of the common garden is outlined in yellow. (B) Breeding design used to generate experimental plants. Field-collected parents were crossed to produce F1 offspring, which were planted in the common garden during the 2008-2009 growing season. These F1s were then selfed to produce F2s, which were planted during the 2009–2010 season. Traits measured in each common garden are listed. (C) Experimental cross types among parental lineages. The three color blocks represent the three mountain regions included in the design. The high-elevation edge population (ME) was situated between the ML and HE transects and served as a key population for inter-range crosses.

**Figure 2.**
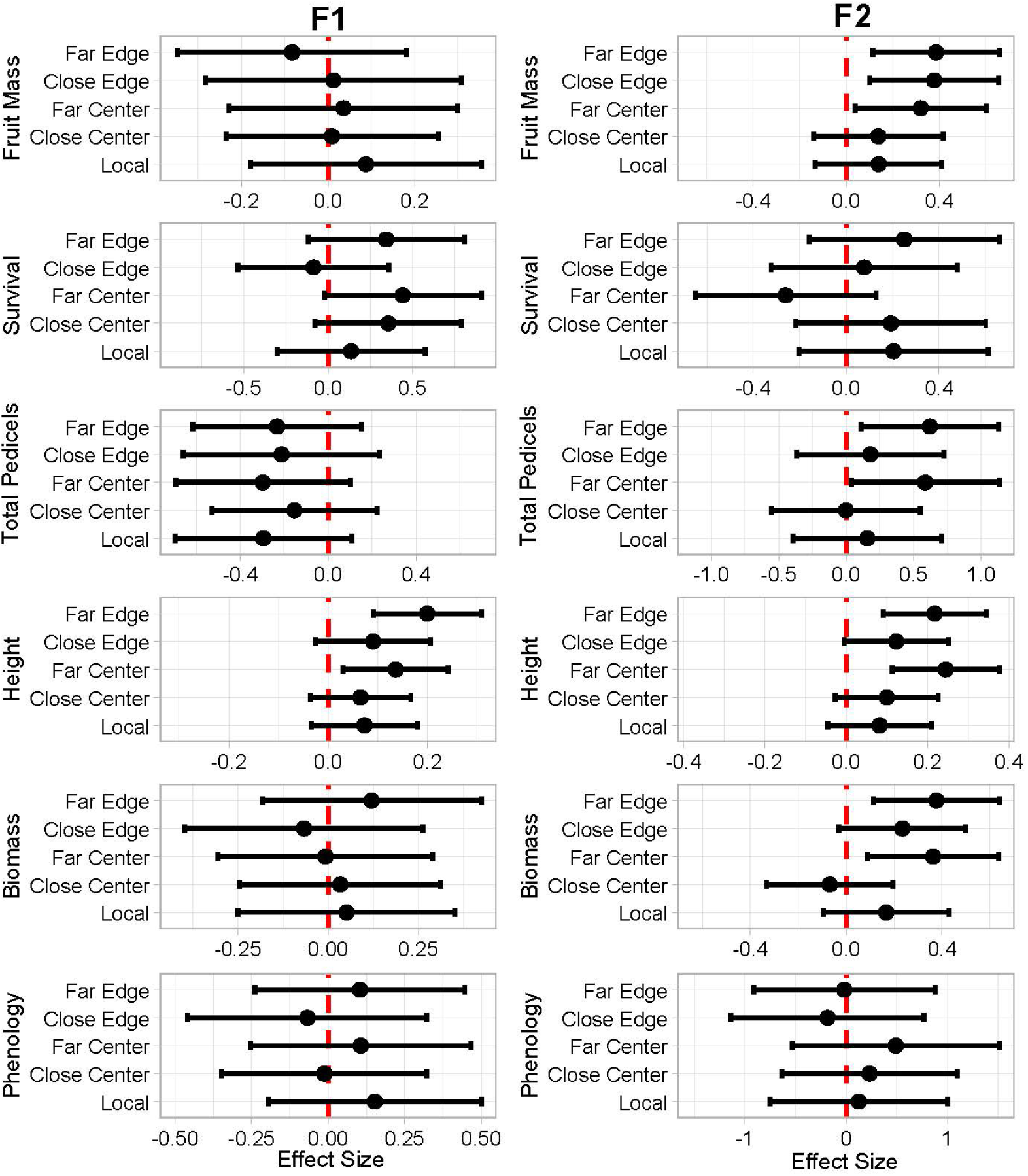
Forest plots showing cross type effects on fitness-related traits in *E. laciniata* relative to self-pollination for 2009 (left panels) and 2010 (right panels). Each panel displays effect sizes (log scale) with 95% confidence intervals for different cross types compared to self-pollination (SELF). Points represent estimated mean differences and horizontal bars show confidence intervals. The red dashed vertical line at zero indicates no difference from self-pollination. Cross types are ordered by distance and location: Local (same population), Close Center and Far Center (central dam locations), and Close Edge and Far Edge (dam edge locations). Effect sizes are derived from generalized linear mixed models with cross type as a fixed effect. Confidence intervals that do not overlap zero indicate statistically significant differences from self-pollination. Traits examined include phenology, biomass, height, total pedicels, survival, and fruit mass across both study years.

**Figure 3.**
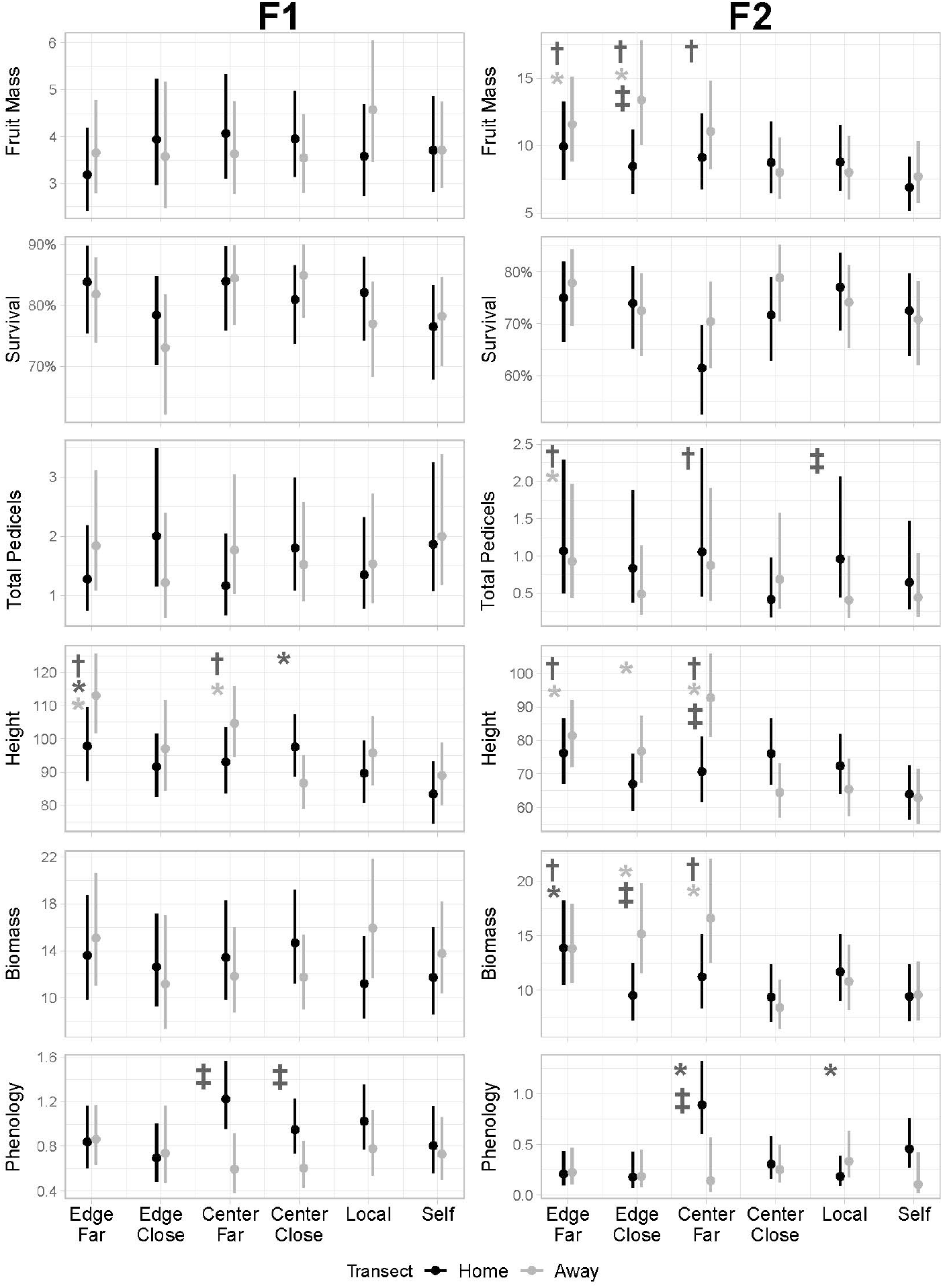
Predicted values from generalized linear mixed models (GLMMs) for final height, total pedicels, phenology, biomass, and lifetime fitness of F1 and F2 generations across cross types and transects (Home, A = black; Away, B = grey). Lifetime fitness was estimated as total fruit mass, including individuals that did not survive to flower (i.e., zero values). Points indicate predicted means for each cross type within each transect, with error bars showing 95% confidence intervals from GLMMTMB models of *[trait] ∼ cross type + transect*. Asterisks denote significant contrasts relative to selfed progeny: † = significant differences between the given cross type (home and away) and selfed progeny; black * = significant differences between home cross types and home selfed progeny; light grey * = significant differences between away cross types and away selfed progeny. ‡ = significant differences between transects for the same trait.

**Figure 4.**
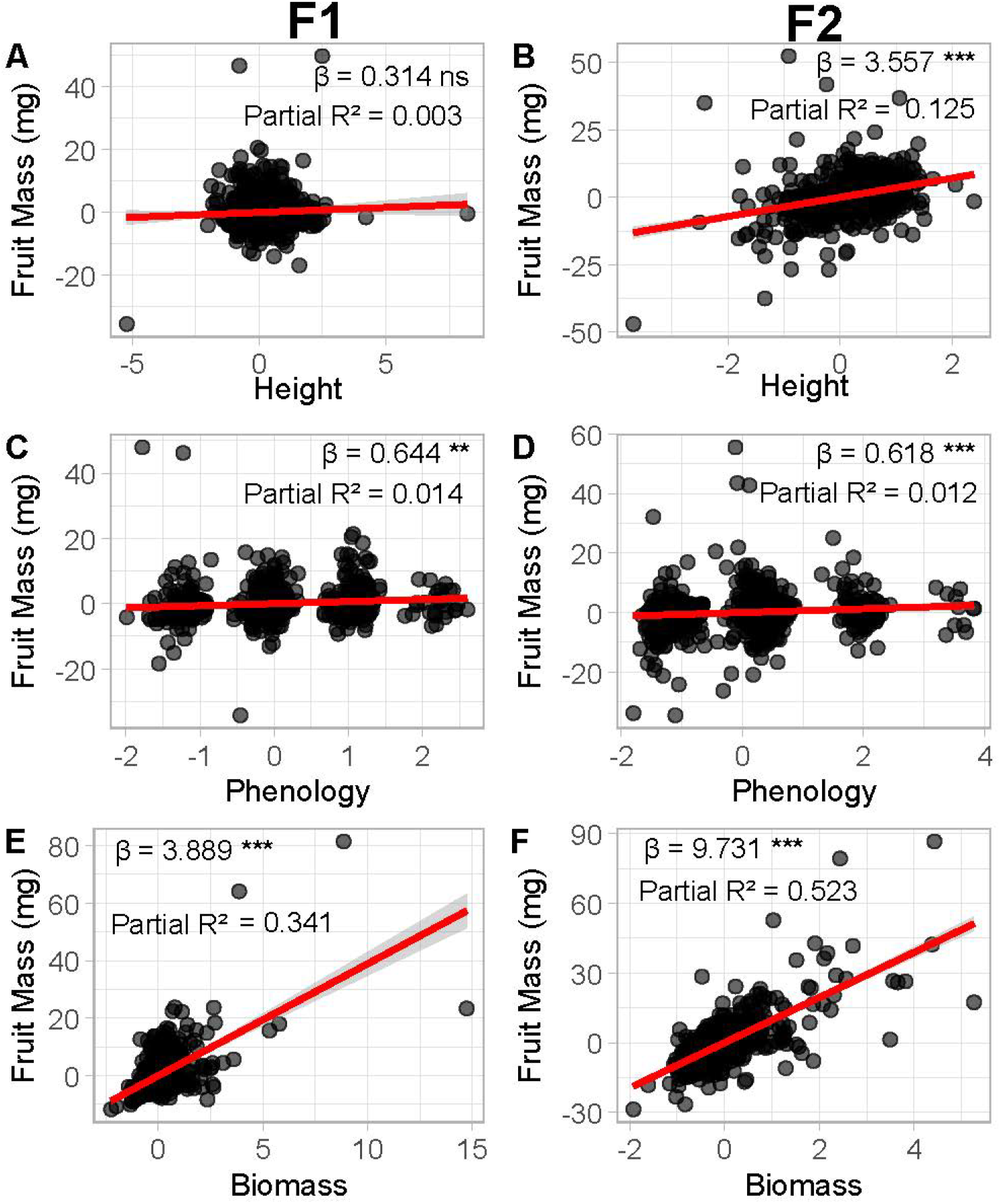
Scatter plots show the relationship between standardized trait values (x-axis) and individual fruit mass (y-axis) for (A, B) final height, (C, D) June phenology, and (E, F) vegetative biomass. Red lines represent linear regression fits with 95% confidence intervals (gray shading). Selection gradients (β) indicate the strength and direction of phenotypic selection, with positive values indicating selection for higher trait values and negative values indicating selection for lower trait values. Significance levels: *** p < 0.001, ** p < 0.01, * p < 0.05, ns = not significant. R^2^ values indicate the proportion of variance in fruit mass explained by each trait. Standardized traits have mean = 0 and standard deviation = 1, allowing for direct comparison of selection strength across different traits and years.

### Experimental Gene Flow Crosses

At least 60 maternal seed families were collected from plants at each population to represent local environmental heterogeneity (see seed collection details in (62)). All plants used for gene flow crosses were the progeny of a refresher generation raised in controlled environment chambers (grown in Sunshine Mix #1 media under 14-h days with 23 °C daytime and 4 °C night-time temperatures) to minimize maternal effects. An F2 generation was created by selfing all F1 offspring to eliminate the influence of hybrid incompatibilities present in the F1 generation, and thus to observe the longer-lasting effects of various gene flow scenarios on reproductive, morphological, and phenological traits (Figure 1). Within each transect, 20 gene-flow recipients (dams) from the upper range limit (subalpine) population received outcrossed pollen originating from one of six sources: (i) 10 randomly selected local sires from the same population (“Local”), (ii) 10 randomly selected sires from a nearby central (lower montane) site on that same transect (“Close Center”), (iii) 10 randomly selected sires from a more distant and lower elevation central site (subalpine) on that same transect (“Far Center”), (iv) 10 randomly selected sires at a high-edge population (subalpine) between transects (“Close Edge”), (v) 10 randomly selected sires from the high-edge population of the other transect (“Far Edge”), and (vi) pollen from the same individual via automatic selfing (“Self”), as a natural control for outcrossing. The same mating design was then repeated for populations in the second transect.

### Common Garden Experiment at the Upper Range Limit

Seeds from experimental crosses (N= 1464 (F1; 2008-2009); N = 1416 (F2; 2009-2010)) were sown randomly into 61 and 59 trays in 2008-2009 and 2009-2010, respectively. Tray cells were filled with a moss-based planting medium that simulates the natural substrate and soil depth *E. laciniata* is adapted to (20, 65). Trays were placed into the field along seeps within naturally growing populations and set onto capillary matting (WA-cpm; Greenhouse Megastore) placed on the rock surfaces. The matting was used to wet the trays via absorption from the wet rock surfaces and surrounding moss of the seeps, and this allowed for natural germination over natural winter conditions while allowing plant roots to expand into the surrounding natural soil matrix. Trays were set in autumn before the onset of the winter season within patches of moss from which grew *E. laciniata* plants the prior year, which was evident from plant skeletons. Five seeds from a single, randomly selected maternal family were sown per position and later thinned to one seedling as an over-sowing approach (as in (20, 65, 78)) to maximize the chance of establishing one independent replicate per maternal family per planting position. Seedlings were later thinned to the most central individual in a cell, early in development before plants were large enough to compete for light or space. Plants were monitored for survival and received a phenological index score (1, vegetative; 2,flower buds present; 3, open flowers present; 4, developing fruit present) referred to here as a “phenological stage.” Plants were harvested at the end of the growing season once all had senesced. Survival to flower, final height, whole plant fruit mass, total fruit mass, and total number of reproductive branches or pedicels (hereafter referred to as “total pedicels”) were also recorded.

### Analysis

Analyses were restricted to blocks that included at least one plant that survived to flowering. Blocks excluded from the analyses (0 out of 61 in 2009 and 2 out of 59 in 2010) were removed due to disturbance (e.g., snowpack movement, animal disruption) or early drying in seep areas that prevented plant development. All analyses were performed separately for the F1 and F2 generation experimental years.

Generalized linear mixed models (GLMMs) were constructed for survival, fruit mass, total pedicels, phenology, final height, and biomass with the interaction of cross type and transect as fixed effects and block as a random effect with an observation-level random effect (OLRE) where appropriate to account for overdispersion. Survival was modeled using a binomial regression with a logit link function. Fruit mass was modelled using a lognormal regression. Phenology and total pedicels were modelled using a zero-inflated truncated negative biominal. Final height and biomass were modelled using a gamma regression with a log link function.

GLMM models were performed with the glmmTMB package in R (79). All models were checked for normality and homoscedasticity of residuals using the simulateResiduals function in the DHARMa R package (v. 0.4.7; (80)), and Type II Wald chi-square tests were used to assess the significance of model terms. For all glmmTMB models, we used the Anova function in the R car package (v.3.1.3; (81)) to determine the significance of the fixed effects in each model. We visualized results using ggplot2 (v.3.5.1; (82)) and ggeffects (v.2.2.1; (83)).

To determine whether outcrossed progeny significantly differed in performance from selfed crosses—the predominant mating strategy of *E. laciniata* (mean FIS = 0.93 across species range, (62)) — pairwise comparisons were conducted using estimated marginal means (EMMs) from the generalized linear mixed models for each trait and generation. For each model, EMMs were computed using the emmeans package in R (84) and pairwise contrasts were extracted using pairs(). Three sets of comparisons were performed from each model: (1) overall contrasts of each cross type against selfed progeny across all samples, (2) contrasts of each cross type against selfed progeny within each source transect (home or away) separately, and (3) contrasts of home versus away source transect within each cross type, to assess whether crosses originating from the local population outperformed those sourced from a geographically distinct population. The absence of a significant omnibus interaction between cross type and transect does not preclude meaningful within-transect differences; the omnibus test evaluates whether the pattern of cross type differences is heterogeneous across transects globally, whereas planned within-transect contrasts detect localized effects for specific cross types that may be consistent in direction but insufficient to drive a significant interaction term. All comparisons were treated as a priori planned contrasts based on explicit biological hypotheses regarding inbreeding depression and local adaptation; therefore, no correction for multiple comparisons was applied. Each comparison includes the estimate, confidence interval, z-ratio, and p-value.

To evaluate fitness outcomes, we used two complementary approaches. First, we applied a Bayesian hurdle model framework that integrates survival and fruit mass into a single composite fitness trait (“lifetime fitness”), thereby capturing the combined probability of surviving and subsequent number of offspring. We fit a series of nested hurdle-gamma models using the brms package in R (85), with block included as a random effect to account for spatial variation in the common garden, including a full model with the interaction between cross type and transect, additive models without interaction, single-predictor models, and an intercept-only null model. Model performance was compared using leave-one-out cross-validation (LOO) and WAIC to assess predictive accuracy across competing models. In addition, we calculated Bayes factors for targeted contrasts, allowing us to quantify the relative contribution for each predictor.

To understand how traits influence fitness variation among individuals at the upper range limit, we performed phenotypic selection analysis (PSA) for phenology, biomass, and height. An individual’s relative fitness can be calculated based on block means or grand means. We found that grand means resulted in a lower AIC score than block means, though both methods produced similar results of significance. We pooled all individuals across all cross types for PSA. This was performed separately for the F1 and F2 generation experimental years.

### Genetic Estimates of Inbreeding and Distance Between Populations

Previous work by Sexton et al. (62) on the same populations extracted DNA using a modified cetyltrimethyl ammonium bromide protocol (86) and used 11 codominant simple sequence repeat (SSR) markers (nuclear-gene-intron-length markers and microsatellites) for population genetic analyses (see TableS4 in (20)). All markers were located on different linkage groups and therefore represented genetically independent loci. Pairwise genetic distance estimates between populations were generated as graph distances using Population Graphs (87) and are published in (62). Genetic distance estimates for populations in this study are given in Table 1.

## Acknowledgements

We thank Joanna Clines, Monique Kolster, Emilio Laca, Bob Latta, and Steve Schoenig for helpful discussions. We thank the following individuals for field and laboratory assistance: Barrett Able, Austin Aslan, Lily Cai, Alexa Carleton, Annie Chang, Donna Chen, Dana Chou, Ruthie Chow, Zacharia Costa, Tomas Gepts, Oscar Gonzalez, Nikhil Gopal, Bryant Gross, Dena Grossenbacher, Yasmine Hernandez, Luke Hoekstra, Carrie Huynh, Christina Islas, Marta Hura, Marcus Jones, Tina Lam, Finn Leeper, Tihua Lee, Jerell Maneja, Lauren McGeoch, Joshua Mopas, Cassandra Morales-de-Silvestore, Thuy Nguyan, Jessenia Perez, Jeff Port, Eli Refsdal, Nerissa Rujanavech, Tam Tran, Randeep Uppal, Marit Wilkerson, and Jennifer Wolf. We acknowledge the Yosemite Field Station (https://doi.org/10.21973/n3v36c) and Valentine Eastern Sierra Reserve (https://doi.org/10.21973/N3966F) in helping conduct this work. Yosemite National Park, Devils Postpile National Monument, the US Forest Service (specifically, Joanna Clines and Diane Ikeda) provided land and plant resources. This work was supported by grants and fellowships (to J.P.S.) from the California Native Plant Society; US Forest Service Native Plant Materials Program NFN3; the National Science Foundation (NSF-DEB 0808607, NSF-IOS-1558035); and the Hellman Fellows Fund at UC Merced.

## References

1. J. H. Brown, On the Relationship between Abundance and Distribution of Species. The American Naturalist 124, 255–279 (1984).

2. R. D. Sagarin, S. D. Gaines, The “Abundant Centre” Distribution: To What Extent Is It a Biogeographical Rule? Ecology Letters 5, 137–147 (2002).

3. R. Hengeveld, J. Haeck, The Distribution of Abundance. I. Measurements. Journal of Biogeography 9, 303 (1982).

4. N. J. Ouborg, P. Vergeer, C. Mix, The Rough Edges of the Conservation Genetics Paradigm for Plants. Journal of Ecology 94, 1233–1248 (2006).

5. C. G. Eckert, K. E. Samis, S. C. Lougheed, Genetic Variation across Species’ Geographical Ranges: The Central–Marginal Hypothesis and Beyond. Molecular Ecology 17, 1170–1188 (2008).

6. S. M. Carlson, C. J. Cunningham, P. A. H. Westley, Evolutionary Rescue in a Changing World. Trends in Ecology & Evolution 29, 521–530 (2014).

7. B. Pujol, J. R. Pannell, Reduced Responses to Selection After Species Range Expansion. Science 321, 96–96 (2008).

8. A. Hampe, R. J. Petit, Conserving biodiversity under climate change: The rear edge matters. Ecology Letters 8, 461–467 (2005).

9. S. Peischl, M. Kirkpatrick, L. Excoffier, Expansion Load and the Evolutionary Dynamics of a Species Range. The American Naturalist 185, E81–E93 (2015).

10. S. Peischl, L. Excoffier, Expansion load: Recessive mutations and the role of standing genetic variation. Molecular Ecology 24, 2084–2094 (2015).

11. Y. Willi, M. Fracassetti, S. Zoller, J. Van Buskirk, Accumulation of Mutational Load at the Edges of a Species Range. Molecular Biology and Evolution 35, 781–791 (2018).

12. P. Lesica, F. W. Allendorf, When Are Peripheral Populations Valuable for Conservation? Conservation Biology 9, 753–760 (1995).

13. M. Slatkin, Gene Flow and the Geographic Structure of Natural Populations. Science 236, 787–792 (1987).

14. R. Frankham, Genetic Rescue of Small Inbred Populations: Meta-analysis Reveals Large and Consistent Benefits of Gene Flow. Molecular Ecology 24, 2610–2618 (2015).

15. P. W. Hedrick, A. Garcia-Dorado, Understanding Inbreeding Depression, Purging, and Genetic Rescue. Trends in Ecology & Evolution 31, 940–952 (2016).

16. D. Tallmon, G. Luikart, R. Waples, The alluring simplicity and complex reality of genetic rescue. Trends in Ecology & Evolution 19, 489–496 (2004).

17. M. Alleaume-benharira, I. R. Pen, O. Ronce, Geographical Patterns of Adaptation within a Species’ Range: Interactions between Drift and Gene Flow. Journal of Evolutionary Biology 19, 203–215 (2006).

18. J. P. Sexton, P. J. McIntyre, A. L. Angert, K. J. Rice, Evolution and Ecology of Species Range Limits. Annual Review of Ecology, Evolution, and Systematics 40, 415–436 (2009).

19. J. Polechová, Is the sky the limit? On the expansion threshold of a species’ range. PLOS Biology 16, e2005372 (2018).

20. J. P. Sexton, S. Y. Strauss, K. J. Rice, Gene flow increases fitness at the warm edge of a species’ range. Proceedings of the National Academy of Sciences 108, 11704–11709 (2011).

21. M. Bontrager, A. L. Angert, Gene flow improves fitness at a range edge under climate change. Evolution Letters 3, 55–68 (2019).

22. Y. Willi, et al., Conservation genetics as a management tool: The five best-supported paradigms to assist the management of threatened species. Proceedings of the National Academy of Sciences 119, e2105076119 (2022).

23. E. C. Bourne, et al., Between Migration Load and Evolutionary Rescue: Dispersal, Adaptation and the Response of Spatially Structured Populations to Environmental Change. Proceedings of the Royal Society B: Biological Sciences 281, 20132795 (2014).

24. P. R. Grant, B. R. Grant, Predicting Microevolutionary Responses to Directional Selection on Heritable Variation. Evolution 49, 241–251 (1995).

25. W. S. Moore, An Evaluation of Narrow Hybrid Zones in Vertebrates. The Quarterly Review of Biology 52, 263–277 (1977).

26. O. Seehausen, Hybridization and adaptive radiation. Trends in Ecology & Evolution 19, 198–207 (2004).

27. D. Charlesworth, B. Charlesworth, Inbreeding Depression and Its Evolutionary Consequences. Annual Review of Ecology and Systematics 18, 237–268 (1987).

28. D. L. Byers, D. M. Waller, Do Plant Populations Purge Their Genetic Load? Effects of Population Size and Mating History on Inbreeding Depression. Annual Review of Ecology and Systematics 30, 479–513 (1999).

29. M. C. Whitlock, P. K. Ingvarsson, T. Hatfield, Local drift load and the heterosis of interconnected populations. Heredity 84, 452–457 (2000).

30. L. H. Rieseberg, M. A. Archer, R. K. Wayne, Transgressive Segregation, Adaptation and Speciation. Heredity 83, 363–372 (1999).

31. D. R. Dittrich-Reed, B. M. Fitzpatrick, Transgressive Hybrids as Hopeful Monsters. Evolutionary Biology 40, 310–315 (2013).

32. R. Stelkens, O. Seehausen, Genetic Distance Between Species Predicts Novel Trait Expression in Their Hybrids. Evolution 63, 884–897 (2009).

33. R. Whitlock, et al., A systematic review of phenotypic responses to between-population outbreeding. Environmental Evidence 2, 13 (2013).

34. J. T. Anderson, M. A. Geber, Demographic Source-Sink Dynamics Restrict Local Adaptation in Elliott’s Blueberry (Vaccinium Elliottii). Evolution 64, 370–384 (2010).

35. T. E. Farkas, T. Mononen, A. A. Comeault, P. Nosil, Observational evidence that maladaptive gene flow reduces patch occupancy in a wild insect metapopulation: BRIEF COMMUNICATION. Evolution 70, 2879–2888 (2016).

36. J. R. Paul, S. N. Sheth, A. L. Angert, Quantifying the Impact of Gene Flow on Phenotype-Environment Mismatch: A Demonstration with the Scarlet Monkeyflower Mimulus Cardinalis. The American Naturalist 178, S62–S79 (2011).

37. M. L. Santon, C. Galen, Life on The Edge: Adaptation Versus Environmentally Mediated Gene Flow in The Snow Buttercup, Ranunculus Adoneus. The American Naturalist 150, 143– 178 (1997).

38. S. Prieto-Benítez, et al., Evaluating Assisted Gene Flow in Marginal Populations of a High Mountain Species. Frontiers in Ecology and Evolution 9, 638837 (2021).

39. J. Morente-López, et al., Gene Flow Effects on Populations Inhabiting Marginal Areas: Origin Matters. Journal of Ecology 109, 139–153 (2021).

40. E. J. Kottler, E. E. Dickman, J. P. Sexton, N. C. Emery, S. J. Franks, Draining the Swamping Hypothesis: Little Evidence That Gene Flow Reduces Fitness at Range Edges. Trends in Ecology & Evolution 36, 533–544 (2021).

41. S. Edmands, Heterosis and Outbreeding Depression in Interpopulation Crosses Spanning a Wide Range of Divergence. Evolution 53, 1757–1768 (1999).

42. A. D. Johansen-Morris, R. G. Latta, Fitness Consequences of Hybridization Between Ecotypes of Avena Barbata: Hybrid Breakdown, Hybrid Vigor, and Transgressive Segregation. Evolution 60, 1585 (2006).

43. J. B. S. Haldane, J. B. S. Haldane, The Relation between Density Regulation and Natural Selection. Proceedings of The Royal Society B: Biological Sciences 145, 306–308 (1956).

44. J. Antonovics, The Input from Population Genetics: “The New Ecological Genetics”. Systematic Botany 1, 233 (1976).

45. M. Kirkpatrick, N. H. Barton, Evolution of a Species’ Range. The American Naturalist 150, 1–23 (1997).

46. T. Lenormand, Gene flow and the limits to natural selection. Trends in Ecology & Evolution 17, 183–189 (2002).

47. S. W. Fitzpatrick, et al., Genomic and Fitness Consequences of Genetic Rescue in Wild Populations. Current Biology 30, 517–522.e5 (2020).

48. D. A. Moeller, M. A. Gebre, Ecological Context of the Evolution of Self-Pollination in Clarkia Xantlana: Poulation Size, Plant Communities, and Reproducttive Assurance. Evolution 59, 786–799 (2005).

49. M. Kirkpatrick, N. H. Barton, Evolution of a Species’ Range. The American Naturalist 150, 1–23 (1997).

50. Z. B. Lippman, D. Zamir, Heterosis: Revisiting the magic. Trends in Genetics 23, 60–66 (2007).

51. D. Charlesworth, J. H. Willis, The genetics of inbreeding depression. Nature Reviews Genetics 10, 783–796 (2009).

52. B. Charlesworth, D. Charlesworth, The genetic basis of inbreeding depression. Genetical Research 74, 329–340 (1999).

53. D. A. Roff, Inbreeding Depression: Tests of the Overdominance and Partial Dominance Hypotheses. Evolution 56, 768–775 (2002).

54. J. K. Kelly, H. S. Arathi, Inbreeding and the genetic variance in floral traits of Mimulus guttatus. Heredity 90, 77–83 (2003).

55. D. E. Carr, M. R. Dudash, Recent approaches into the genetic basis of inbreeding depression in plants. Philosophical Transactions of the Royal Society of London. Series B: Biological Sciences 358, 1071–1084 (2003).

56. M. Nei, The frequency distribution of lethal chromosomes in finite populations. Proceedings of the National Academy of Sciences 60, 517–524 (1968).

57. S. Glémin, J. Ronfort, T. Bataillon, Patterns of Inbreeding Depression and Architecture of the Load in Subdivided Populations. Genetics 165, 2193–2212 (2003).

58. J. F. Crow, “Genetic Loads and the Cost of Natural Selection“ in Mathematical Topics in Population Genetics, K. Kojima, Ed. (Springer Berlin Heidelberg, 1970), pp. 128–177.

59. J. F. Crow, M. Kimura, An Introduction to Population Genetics Theory. Population (French Edition) 26, 977 (1971).

60. A. García-Fernández, J. M. Iriondo, A. Escudero, Inbreeding at the Edge: Does Inbreeding Depression Increase under More Stressful Conditions? Oikos 121, 1435–1445 (2012).

61. D. Roze, F. Rousset, Joint Effects of Self-Fertilization and Population Structure on Mutation Load, Inbreeding Depression and Heterosis. Genetics 167, 1001–1015 (2004).

62. J. P. Sexton, et al., Climate structures genetic variation across a species’ elevation range: A test of range limits hypotheses. Molecular Ecology 25, 911–928 (2016).

63. G. García-Ramos, M. Kirkpatrick, Genetic Models of Adaptation and Gene Flow in Peripheral Populations. Evolution 51, 21–28 (1997).

64. J. Polechová, N. H. Barton, Limits to Adaptation along Environmental Gradients. Proceedings of the National Academy of Sciences 112, 6401–6406 (2015).

65. J. E. Shay, L. K. Pennington, D. J. Toews, E. Green, J. P. Sexton, The Leading Edge Matters Too: Fitness and the Expression of Adaptive Differentiation Are Greatest at the High-Elevation Edge of a Species’ Range. Ecology Letters 29, e70329 (2026).

66. E. P. Kingsley, M. Manceau, C. D. Wiley, H. E. Hoekstra, Melanism in Peromyscus Is Caused by Independent Mutations in Agouti. PLoS ONE 4, e6435 (2009).

67. R. Greenway, et al., Convergent Evolution of Conserved Mitochondrial Pathways Underlies Repeated Adaptation to Extreme Environments. (2020).

68. S. Chaturvedi, et al., Climatic Similarity and Genomic Background Shape the Extent of Parallel Adaptation in Timema Stick Insects. Nature Ecology and Evolution (2022). 10.1038/s41559-022-01909-6.

69. D. M. Weinreich, R. A. Watson, L. Chao, Perspective: Sign Epistasis and Genetic Costraint on Evolutionary Trajectories. Evolution 59, 1165–1174 (2005).

70. L. Ferretti, D. Weinreich, F. Tajima, G. Achaz, Evolutionary constraints in fitness landscapes. Heredity 121, 466–481 (2018).

71. A. H. Patton, E. J. Richards, K. J. Gould, L. K. Buie, C. H. Martin, Hybridization alters the shape of the genotypic fitness landscape, increasing access to novel fitness peaks during adaptive radiation. eLife 11, e72905 (2022).

72. S. N. Aitken, M. C. Whitlock, Assisted Gene Flow to Facilitate Local Adaptation to Climate Change. Annual Review of Ecology, Evolution, and Systematics 44, 367–388 (2013).

73. J. P. Sexton, et al., Patterns and effects of gene flow on adaptation across spatial scales: Implications for management. Journal of Evolutionary Biology 37, 732–745 (2024).

74. S. Mezmouk, J. Ross-Ibarra, The Pattern and Distribution of Deleterious Mutations in Maize. G3 GenesGenomesGenetics 4, 163–171 (2014).

75. B. M. Moran, et al., The genomic consequences of hybridization. eLife 10, e69016 (2021).

76. M. Hasselgren, et al., Strongly deleterious mutations influence reproductive output and longevity in an endangered population. Nature Communications 15, 8378 (2024).

77. E. Hart, K. Bell, Prism: Access data from the Oregon State Prism climate project. (2015).

78. E. E. Dickman, L. K. Pennington, S. J. Franks, J. P. Sexton, Evidence for adaptive responses to historic drought across a native plant species range. Evolutionary Applications 12, 1569–1582 (2019).

79. M. Brooks E., et al., glmmTMB Balances Speed and Flexibility Among Packages for Zero-inflated Generalized Linear Mixed Modeling. The R Journal 9, 378 (2017).

80. F. Hartig, DHARMa: Residual Diagnostics for Hierarchical (Multi-Level / Mixed) Regression Models. 0.4.7 (2016).

81. J. Fox, S. Weisberg, An R companion to applied regression, 2nd edition (Sage, 2011).

82. H. Wickham, Ggplot2 (Springer International Publishing, 2016).

83. D. Lüdecke, Ggeffects: Tidy Data Frames of Marginal Effects from Regression Models. Journal of Open Source Software 3, 772 (2018).

84. R. V. Lenth, J. Piaskowski, Emmeans: Estimated Marginal Means, aka Least-Squares Means. 2.0.1 (2017).

85. P.-C. Bürkner, Brms : An R Package for Bayesian Multilevel Models Using Stan. Journal of Statistical Software 80 (2017).

86. J.-Z. Lin, K. Ritland, Flower petals allow simpler and better isolation of DNA for plant RAPD analyses. Plant Molecular Biology Reporter 13, 210–213 (1995).

87. R. J. Dyer, GeneticStudio: A suite of programs for spatial analysis of genetic-marker data. Molecular Ecology Resources 9, 110–113 (2009).

