## Supplemental Figures and Tables for "Greater benefits of assisted gene flow in F2 vs F1 progeny at the cold edge of a species’ range"

Supplemental Table 1. Average mean, maximum, and minimum temperature, including total precipitation, of each site experimental populations originated from for 2008-2009 (F1), 2009-2010 (F2), and a thirty-year baseline from 1981-2010. Daily measures were extracted for each site's latitude and longitude using the prism package in R (Hart & Bell 2015).

| Climate Variable | Time Period | BC | BP | HE | ME | ML | SC1 | TN |
|---|---|---|---|---|---|---|---|---|
| Mean | 2008-2009 | 1.39 | 7.80 | 1.39 | 4.22 | 4.55 | 7.31 | 5.55 |
| Temperature | 2009-2010 | 0.50 | 6.33 | 0.50 | 3.22 | 3.70 | 6.38 | 4.46 |
| | 1981-2010 | 1.18 | 7.11 | 1.18 | 3.21 | 3.84 | 6.53 | 4.38 |
| Max | 2008-2009 | 7.73 | 13.37 | 7.73 | 10.41 | 11.65 | 13.98 | 12.19 |
| Temperature | 2009-2010 | 7.10 | 11.98 | 7.10 | 9.31 | 10.74 | 13.01 | 11.05 |
| | 1981-2010 | 7.62 | 12.8 | 7.62 | 9.86 | 10.9 | 13.2 | 11.5 |
| Min | 2008-2009 | -4.94 | 2.23 | -4.94 | -1.97 | -2.56 | 0.64 | -1.09 |
| Temperature | 2009-2010 | -6.10 | 0.69 | -6.10 | -2.88 | -3.34 | -0.25 | -2.14 |
| | 1981-2010 | -5.27 | 1.40 | -5.27 | -3.44 | -3.22 | -0.11 | -2.73 |
| Total | 2008-2009 | 1219 | 972 | 1219 | 822 | 1296 | 1114 | 1156 |
| Precipitation | 2009-2010 | 1388 | 1228 | 1388 | 769 | 1552 | 1310 | 1377 |
| | 1981-2010 | 1163 | 1178 | 1164 | 787 | 1458 | 1241 | 1273 |

Supplemental Table 2. Pairwise comparisons of each cross type to the control selfed progeny for F1 and F2 generations grown at *E. laciniatus'* high elevation range edge. Each row shows the 95% confidence interval, z-ratio, and p-value for each contrast and trait. Results are based on estimated marginal means (emmeans) from generalized linear mixed models.

| | | | F1 | | | F2 | |
|---|---|---|---|---|---|---|---|
| Trait | Contrasts | CI | z | p | CI | z | p |
| Fruit Mass | Close Center vs Self | -0.236-0.254 | 0.071 | 0.943 | -0.14-0.417 | 0.973 | 0.33 |
| | Far Center vs Self | -0.229-0.299 | 0.261 | 0.794 | 0.038-0.603 | 2.227 | 0.026 |
| | Close Edge vs Self | -0.283-0.307 | 0.080 | 0.936 | 0.101-0.656 | 2.67 | 0.008 |
| | Far Edge vs Self | -0.348-0.181 | -0.616 | 0.538 | 0.115-0.658 | 2.789 | 0.005 |
| | Local vs Self | -0.179-0.353 | 0.642 | 0.521 | -0.133-0.412 | 1.005 | 0.315 |
| Survival | Close Center vs Self | -0.077-0.79 | 1.61 | 0.107 | -0.216-0.601 | 0.923 | 0.356 |
| | Far Center vs Self | -0.023-0.906 | 1.864 | 0.062 | -0.651-0.13 | -1.309 | 0.19 |
| | Close Edge vs Self | -0.533-0.361 | -0.379 | 0.705 | -0.323-0.479 | 0.381 | 0.703 |
| | Far Edge vs Self | -0.117-0.805 | 1.462 | 0.144 | -0.159-0.659 | 1.198 | 0.231 |
| | Local vs Self | -0.304-0.575 | 0.606 | 0.545 | -0.204-0.612 | 0.979 | 0.328 |
| Total Pedicels | Close Center vs Self | -0.529-0.222 | -0.802 | 0.423 | -0.551-0.549 | -0.005 | 0.996 |
| | Far Center vs Self | -0.695-0.1 | -1.465 | 0.143 | 0.038-1.134 | 2.096 | 0.036 |
| | Close Edge vs Self | -0.659-0.234 | -0.932 | 0.351 | -0.365-0.723 | 0.644 | 0.519 |
| | Far Edge vs Self | -0.617-0.152 | -1.185 | 0.236 | 0.112-1.131 | 2.389 | 0.017 |
| | Local vs Self | -0.696-0.107 | -1.439 | 0.15 | -0.394-0.707 | 0.557 | 0.578 |
| Height | Close Center vs Self | -0.036-0.166 | 1.267 | 0.205 | -0.027-0.227 | 1.546 | 0.122 |
| | Far Center vs Self | 0.03-0.242 | 2.511 | 0.012 | 0.113-0.376 | 3.649 | <0.0001 |
| | Close Edge vs Self | -0.025-0.206 | 1.536 | 0.125 | -0.004-0.251 | 1.894 | 0.058 |
| | Far Edge vs Self | 0.091-0.308 | 3.595 | <0.0001 | 0.092-0.343 | 3.386 | 0.001 |
| | Local vs Self | -0.034-0.18 | 1.338 | 0.181 | -0.044-0.209 | 1.272 | 0.203 |
| Biomass | Close Center vs Self | -0.245-0.312 | 0.236 | 0.813 | -0.332-0.195 | -0.508 | 0.612 |
| | Far Center vs Self | -0.305-0.29 | -0.049 | 0.961 | 0.09-0.636 | 2.603 | 0.009 |
| | Close Edge vs Self | -0.397-0.263 | -0.397 | 0.691 | -0.03-0.499 | 1.738 | 0.082 |
| | Far Edge vs Self | -0.183-0.424 | 0.778 | 0.436 | 0.115-0.639 | 2.821 | 0.005 |
| | Local vs Self | -0.249-0.351 | 0.333 | 0.739 | -0.095-0.429 | 1.25 | 0.211 |
| Phenology | Close Center vs Self | -0.348-0.323 | -0.075 | 0.94 | -0.637-1.099 | 0.522 | 0.602 |
| | Far Center vs Self | -0.253-0.467 | 0.583 | 0.56 | -0.532-1.511 | 0.939 | 0.348 |
| | Close Edge vs Self | -0.458-0.323 | -0.34 | 0.734 | -1.138-0.769 | -0.379 | 0.704 |
| | Far Edge vs Self | -0.238-0.446 | 0.595 | 0.552 | -0.913-0.877 | -0.039 | 0.969 |
| | Local vs Self | -0.195-0.5 | 0.859 | 0.39 | -0.754-1.002 | 0.277 | 0.781 |

Supplemental Table 3. Bayesian model comparison using Watanabe-Akaike Information Criterion (WAIC) for hurdle lognormal models predicting composite fitness in *E. laciniata* for 2009 and 2010. Models are ranked from best (lowest WAIC) to worst (highest WAIC). The elpd_diff column shows the difference in expected log pointwise predictive density relative to the best model (set to 0.0), with negative values indicating worse predictive performance. The se_diff column shows the standard error of the difference. Models within 2 WAIC units are considered to have equivalent predictive performance. Model specifications: null = intercept-only; no dam = cross type only; no cross = dam transect only; no int = additive effects of cross type and dam transect; full = interaction between cross type and dam transect. In 2009, the null model performed best, suggesting limited evidence for treatment effects. In 2010, the cross type-only model had the best predictive performance, followed closely by the null and additive models (differences < 1 WAIC unit), indicating weak but potentially meaningful cross type effects.

| F1 | elpd_diff | se_diff |
|---|---|---|
| ~ Cross Type + Transect + Cross Type * Transect | -15.3 | 4 |
| ~ Cross Type + Transect | -7.4 | 3.2 |
| ~Cross Type | -2.1 | 0.2 |
| ~Transect | -5.3 | 3.3 |
| ~ | 0 | 0 |
| F2 | elpd_diff | se_diff |
| ~ Cross Type + Transect + Cross Type * Transect | -5.3 | 3.5 |
| ~ Cross Type + Transect | -0.4 | 1.6 |
| ~Cross Type | -0.9 | 4.8 |
| ~Transect | 0 | 0 |
| ~ | -0.4 | 4.5 |

Supplemental Table 4. Pairwise comparisons of each cross type by transect of F1 and F2 generations grown at *E. laciniatus'* high elevation range edge. Each row shows the 95% confidence interval, z-ratio, and p-value for each contrast and trait. Results are based on estimated marginal means (emmeans) from generalized linear mixed models.

| | | | F1 | | | F2 | |
|---|---|---|---|---|---|---|---|
| Model | Cross Type | CI | z | p | CI | z | p |
| Fruit Mass | Close Center | -0.218-0.436 | 0.653 | 0.514 | -0.309-0.483 | 0.43 | 0.667 |
| | Far Center | -0.268-0.494 | 0.583 | 0.56 | -0.598-0.214 | -0.927 | 0.354 |
| | Close Edge | -0.368-0.56 | 0.407 | 0.684 | -0.849--0.07 | -2.31 | 0.021 |
| | Far Edge | -0.52-0.246 | -0.701 | 0.483 | -0.529-0.225 | -0.789 | 0.43 |
| | Local | -0.631-0.143 | -1.234 | 0.217 | -0.289-0.47 | 0.468 | 0.639 |
| | Self | -0.366-0.364 | -0.006 | 0.995 | -0.504-0.279 | -0.561 | 0.575 |
| Survival | Close Center | -0.9-0.341 | -0.882 | 0.378 | -0.98-0.208 | -1.273 | 0.203 |
| | Far Center | -0.74-0.665 | -0.104 | 0.917 | -0.942-0.141 | -1.45 | 0.147 |
| | Close Edge | -0.367-0.948 | 0.867 | 0.386 | -0.499-0.647 | 0.253 | 0.8 |
| | Far Edge | -0.555-0.835 | 0.395 | 0.693 | -0.754-0.435 | -0.526 | 0.599 |
| | Local | -0.32-0.952 | 0.975 | 0.33 | -0.434-0.75 | 0.523 | 0.601 |
| | Self | -0.704-0.509 | -0.315 | 0.753 | -0.48-0.644 | 0.286 | 0.775 |
| Total Pedicels | Close Center | -0.335-0.671 | 0.656 | 0.512 | -1.282-0.282 | -1.252 | 0.21 |
| | Far Center | -0.983-0.152 | -1.435 | 0.151 | -0.584-0.962 | 0.479 | 0.632 |
| | Close Edge | -0.199-1.198 | 1.402 | 0.161 | -0.231-1.301 | 1.368 | 0.171 |
| | Far Edge | -0.898-0.163 | -1.358 | 0.175 | -0.518-0.798 | 0.417 | 0.677 |
| | Local | -0.709-0.448 | -0.442 | 0.658 | 0.068-1.639 | 2.13 | 0.033 |
| | Self | -0.626-0.488 | -0.243 | 0.808 | -0.4-1.148 | 0.947 | 0.344 |
| Height | Close Center | -0.014-0.25 | 1.752 | 0.08 | -0.013-0.343 | 1.816 | 0.069 |
| | Far Center | -0.265-0.03 | -1.566 | 0.117 | -0.462--0.08 | -2.78 | 0.005 |
| | Close Edge | -0.231-0.115 | -0.657 | 0.511 | -0.316-0.045 | -1.472 | 0.141 |
| | Far Edge | -0.298-0.01 | -1.832 | 0.067 | -0.242-0.109 | -0.739 | 0.46 |
| | Local | -0.216-0.084 | -0.864 | 0.387 | -0.076-0.279 | 1.123 | 0.261 |
| | Self | -0.218-0.088 | -0.836 | 0.403 | -0.163-0.198 | 0.19 | 0.85 |
| Biomass | Close Center | -0.152-0.593 | 1.158 | 0.247 | -0.262-0.479 | 0.573 | 0.566 |
| | Far Center | -0.301-0.552 | 0.577 | 0.564 | -0.788-0.008 | -1.919 | 0.055 |
| | Close Edge | -0.391-0.635 | 0.465 | 0.642 | -0.838--0.092 | -2.442 | 0.015 |
| | Far Edge | -0.542-0.335 | -0.462 | 0.644 | -0.363-0.369 | 0.017 | 0.987 |
| | Local | -0.785-0.08 | -1.596 | 0.111 | -0.287-0.446 | 0.425 | 0.671 |
| | Self | -0.573-0.255 | -0.753 | 0.452 | -0.39-0.359 | -0.082 | 0.934 |
| Phenology | Close Center | 0.024-0.87 | 2.071 | 0.038 | -0.737-1.127 | 0.409 | 0.682 |
| | Far Center | 0.223-1.215 | 2.84 | 0.005 | 0.394-3.245 | 2.502 | 0.012 |
| | Close Edge | -0.642-0.522 | -0.202 | 0.84 | -1.258-1.185 | -0.059 | 0.953 |
| | Far Edge | -0.474-0.415 | -0.129 | 0.897 | -1.107-0.953 | -0.147 | 0.883 |
| | Local | -0.19-0.73 | 1.152 | 0.249 | -1.555-0.382 | -1.186 | 0.235 |
| | Self | -0.425-0.617 | 0.363 | 0.717 | -0.006-2.922 | 1.952 | 0.051 |

Supplemental Table 5. Pairwise comparisons of each cross type to the control selfed progeny by transect for F1 and F2 generations grown at *E. laciniatus'* high elevation range edge. Each row shows the 95% confidence interval, z-ratio, and p-value for each contrast and trait. Results are based on estimated marginal means (emmeans) from generalized linear mixed models. Home transect = Transect from which high elevation edge population originated. Away transect= transect spanning similar elevations on a different watershed.

| | | | | F1 | | | F2 | |
|---|---|---|---|---|---|---|---|---|
| Model | Transect | Contrast | CI | z | p | CI | z | p |
| Fruit Mass | Home | Close Center vs Self | -0.292-0.42 | 0.352 | 0.725 | -0.156-0.632 | 1.182 | 0.237 |
| | Home | Far Center vs Self | -0.291-0.475 | 0.473 | 0.636 | -0.119-0.681 | 1.375 | 0.169 |
| | Home | Close Edge vs Self | -0.33-0.452 | 0.305 | 0.761 | -0.181-0.59 | 1.041 | 0.298 |
| | Home | Far Edge vs Self | -0.536-0.234 | -0.77 | 0.442 | -0.017-0.75 | 1.872 | 0.061 |
| | Home | Local vs Self | -0.415-0.347 | -0.176 | 0.861 | -0.137-0.619 | 1.25 | 0.211 |
| | Away | Close Center vs Self | -0.384-0.292 | -0.268 | 0.789 | -0.354-0.432 | 0.193 | 0.847 |
| | Away | Far Center vs Self | -0.385-0.341 | -0.119 | 0.906 | -0.037-0.759 | 1.775 | 0.076 |
| | Away | Close Edge vs Self | -0.479-0.406 | -0.162 | 0.871 | 0.154-0.95 | 2.72 | 0.007 |
| | Away | Far Edge vs Self | -0.379-0.348 | -0.082 | 0.934 | 0.022-0.791 | 2.071 | 0.038 |
| | Away | Local vs Self | -0.163-0.58 | 1.099 | 0.272 | -0.354-0.431 | 0.191 | 0.848 |
| Survival | Home | Close Center vs Self | -0.331-0.862 | 0.873 | 0.383 | -0.606-0.523 | -0.144 | 0.886 |
| | Home | Far Center vs Self | -0.192-1.135 | 1.393 | 0.164 | -1.044-0.04 | -1.816 | 0.069 |
| | Home | Close Edge vs Self | -0.499-0.714 | 0.348 | 0.728 | -0.499-0.647 | 0.253 | 0.8 |
| | Home | Far Edge vs Self | -0.212-1.138 | 1.343 | 0.179 | -0.446-0.705 | 0.44 | 0.66 |
| | Home | Local vs Self | -0.289-0.974 | 1.064 | 0.288 | -0.34-0.824 | 0.814 | 0.416 |
| | Away | Close Center vs Self | -0.183-1.078 | 1.391 | 0.164 | -0.165-1.018 | 1.413 | 0.158 |
| | Away | Far Center vs Self | -0.238-1.061 | 1.242 | 0.214 | -0.581-0.542 | -0.067 | 0.947 |
| | Away | Close Edge vs Self | -0.938-0.377 | -0.836 | 0.403 | -0.48-0.644 | 0.286 | 0.775 |
| | Away | Far Edge vs Self | -0.404-0.854 | 0.702 | 0.483 | -0.21-0.952 | 1.25 | 0.211 |
| | Away | Local vs Self | -0.682-0.54 | -0.228 | 0.82 | -0.407-0.738 | 0.568 | 0.57 |
| Total Pedicels | Home | Close Center vs Self | -0.573-0.503 | -0.127 | 0.899 | -1.212-0.336 | -1.11 | 0.267 |
| | Home | Far Center vs Self | -1.047-0.106 | -1.6 | 0.11 | -0.31-1.297 | 1.203 | 0.229 |
| | Home | Close Edge vs Self | -0.513-0.656 | 0.241 | 0.809 | -0.5-1.019 | 0.669 | 0.503 |
| | Home | Far Edge vs Self | -0.94-0.177 | -1.339 | 0.181 | -0.221-1.229 | 1.363 | 0.173 |
| | Home | Local vs Self | -0.891-0.24 | -1.128 | 0.26 | -0.328-1.12 | 1.073 | 0.283 |
| | Away | Close Center vs Self | -0.796-0.251 | -1.019 | 0.308 | -0.347-1.218 | 1.092 | 0.275 |
| | Away | Far Center vs Self | -0.672-0.423 | -0.444 | 0.657 | -0.065-1.422 | 1.789 | 0.074 |
| | Away | Close Edge vs Self | -1.172-0.179 | -1.442 | 0.149 | -0.681-0.879 | 0.248 | 0.804 |
| | Away | Far Edge vs Self | -0.612-0.445 | -0.309 | 0.758 | 0.024-1.452 | 2.026 | 0.043 |
| | Away | Local vs Self | -0.834-0.306 | -0.908 | 0.364 | -0.914-0.748 | -0.196 | 0.844 |
| Height | Home | Close Center vs Self | 0.011-0.303 | 2.101 | 0.036 | -0.006-0.353 | 1.898 | 0.058 |
| | Home | Far Center vs Self | -0.045-0.264 | 1.392 | 0.164 | -0.086-0.287 | 1.055 | 0.291 |
| | Home | Close Edge vs Self | -0.058-0.246 | 1.215 | 0.225 | -0.132-0.225 | 0.514 | 0.607 |

| | | | | | | | | |
|---|---|---|---|---|---|---|---|---|
| | Home | Far Edge vs Self | 0.001-0.318 | 1.975 | 0.048 | -0.003-0.354 | 1.93 | 0.054 |
| | Home | Local vs Self | -0.08-0.225 | 0.936 | 0.349 | -0.051-0.3 | 1.388 | 0.165 |
| | Away | Close Center vs Self | -0.165-0.113 | -0.37 | 0.712 | -0.153-0.205 | 0.287 | 0.774 |
| | Away | Far Center vs Self | 0.017-0.308 | 2.184 | 0.029 | 0.204-0.574 | 4.115 | <0.001 |
| | Away | Close Edge vs Self | -0.087-0.261 | 0.979 | 0.328 | 0.017-0.382 | 2.145 | 0.032 |
| | Away | Far Edge vs Self | 0.09-0.387 | 3.151 | 0.002 | 0.082-0.437 | 2.86 | 0.004 |
| | Away | Local vs Self | -0.077-0.224 | 0.957 | 0.339 | -0.142-0.222 | 0.43 | 0.667 |
| Biomass | Home | Close Center vs Self | -0.181-0.628 | 1.081 | 0.279 | -0.38-0.368 | -0.033 | 0.974 |
| | Home | Far Center vs Self | -0.299-0.569 | 0.609 | 0.543 | -0.213-0.565 | 0.887 | 0.375 |
| | Home | Close Edge vs Self | -0.358-0.505 | 0.333 | 0.739 | -0.361-0.38 | 0.052 | 0.958 |
| | Home | Far Edge vs Self | -0.292-0.588 | 0.66 | 0.509 | 0.014-0.758 | 2.033 | 0.042 |
| | Home | Local vs Self | -0.477-0.386 | -0.207 | 0.836 | -0.149-0.579 | 1.157 | 0.247 |
| | Away | Close Center vs Self | -0.539-0.227 | -0.798 | 0.425 | -0.501-0.241 | -0.688 | 0.491 |
| | Away | Far Center vs Self | -0.556-0.257 | -0.722 | 0.47 | 0.166-0.934 | 2.805 | 0.005 |
| | Away | Close Edge vs Self | -0.706-0.292 | -0.814 | 0.416 | 0.082-0.836 | 2.386 | 0.017 |
| | Away | Far Edge vs Self | -0.323-0.508 | 0.437 | 0.662 | -0.001-0.736 | 1.955 | 0.051 |
| | Away | Local vs Self | -0.269-0.564 | 0.695 | 0.487 | -0.258-0.497 | 0.621 | 0.534 |
| Phenology | Home | Close Center vs Self | -0.282-0.607 | 0.716 | 0.474 | -1.215-0.414 | -0.964 | 0.335 |
| | Home | Far Center vs Self | -0.021-0.857 | 1.866 | 0.062 | 0.028-1.312 | 2.047 | 0.041 |
| | Home | Close Edge vs Self | -0.662-0.371 | -0.554 | 0.58 | -1.933-0.07 | -1.824 | 0.068 |
| | Home | Far Edge vs Self | -0.449-0.531 | 0.165 | 0.869 | -1.673-0.102 | -1.735 | 0.083 |
| | Home | Local vs Self | -0.22-0.699 | 1.021 | 0.307 | -1.786--0.01 | -1.981 | 0.048 |
| | Away | Close Center vs Self | -0.691-0.314 | -0.734 | 0.463 | -0.67-2.396 | 1.103 | 0.27 |
| | Away | Far Center vs Self | -0.774-0.366 | -0.702 | 0.482 | -1.631-2.249 | 0.312 | 0.755 |
| | Away | Close Edge vs Self | -0.575-0.596 | 0.035 | 0.972 | -1.06-2.186 | 0.68 | 0.497 |
| | Away | Far Edge vs Self | -0.312-0.645 | 0.683 | 0.494 | -0.805-2.305 | 0.946 | 0.344 |
| | Away | Local vs Self | -0.456-0.587 | 0.246 | 0.806 | -0.368-2.661 | 1.484 | 0.138 |

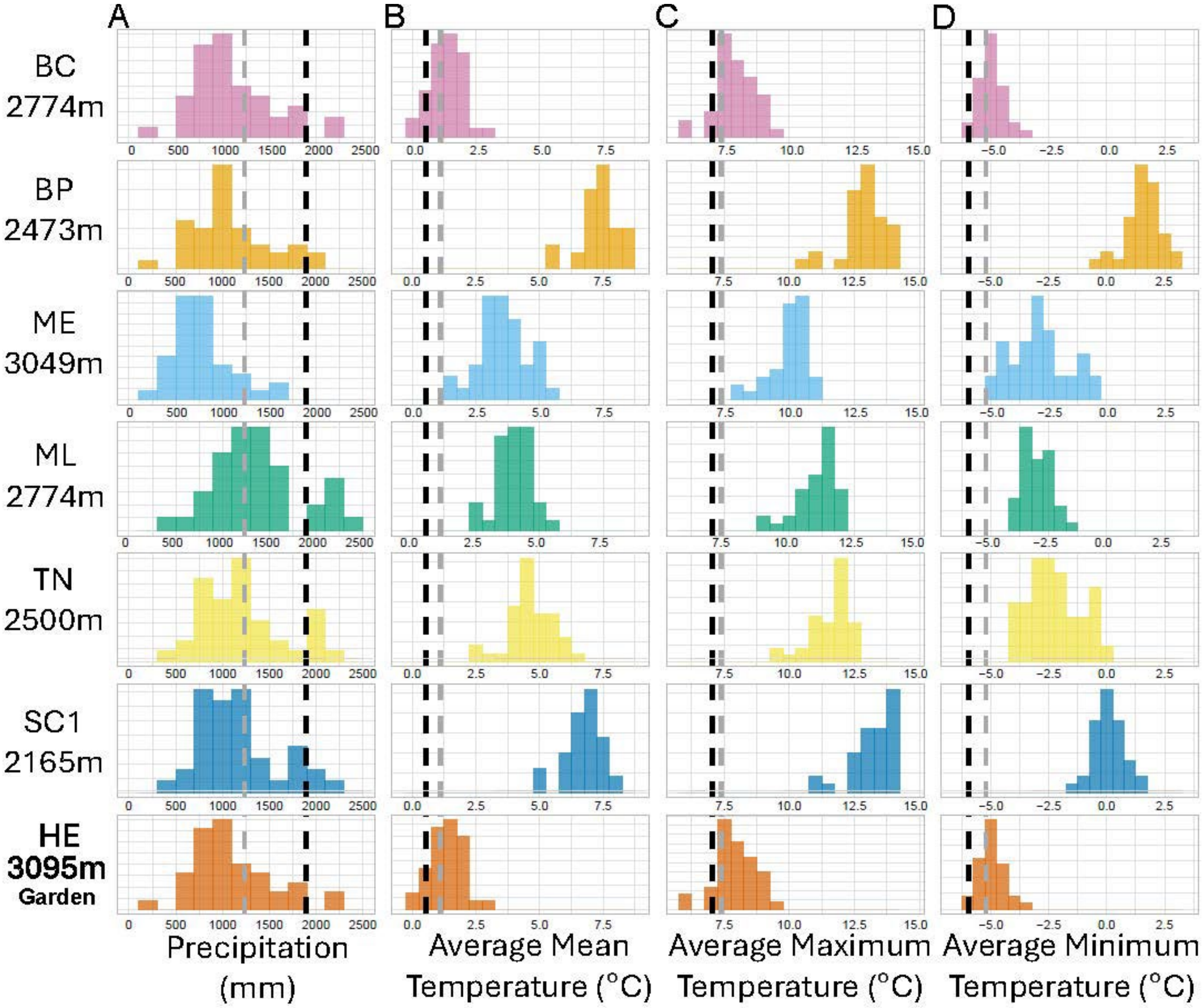

Supplemental Figure 1. Histograms of average (A) accumulated precipitation (B) mean temperature (C) maximum temperature and (D) and minimum temperature from 1981-2010 Dashed lines indicate the climate year of 2009 (grey) and 2010 (black) at the common garden site (HE).
